# Does the phase of ongoing EEG oscillations predict auditory perception?

**DOI:** 10.1101/2020.07.17.209387

**Authors:** I. Tal, M. Leszczynski, N. Mesgarani, C.E. Schroeder

**Affiliations:** Department of Psychiatry; Columbia University Medical Center; New York, NY, 10032; USA; Translational Neuroscience Division of the Center for Biomedical Imaging and Neuromodulation; Nathan Kline Institute for Psychiatric Research; Orangeburg, NY, 10962; USA; Department of Electrical Engineering; Columbia University; New York, NY, 10027; USA

**Keywords:** EEG, oscillations, phase, auditory discrimination

## Abstract

Effective processing of information from the environment requires the brain to selectively sample relevant inputs. The visual perceptual system has been shown to sample information rhythmically, oscillating rapidly between more and less input-favorable states. Evidence of parallel effects in auditory perception is inconclusive. Here, we combined a bilateral pitch-identification task with electroencephalography (EEG) to investigate whether the phase of ongoing EEG predicts auditory discrimination accuracy. We compared prestimulus phase distributions between correct and incorrect trials. Shortly before stimulus onset, each of these distributions showed significant phase concentration, but centered at different phase angles. The effects were strongest in theta and beta frequency bands. The divergence between phase distributions showed a linear relation with accuracy, accounting for at least 10% of inter-individual variance. Discrimination performance oscillated rhythmically at a rate predicted by the neural data. These findings indicate that auditory discrimination threshold oscillates over time along with the phase of ongoing EEG activity. Thus, it appears that auditory perception is discrete rather than continuous, with the phase of ongoing EEG oscillations shaping auditory perception by providing a temporal reference frame for information processing.

## Introduction

Fluctuations of excitability in ensembles of neurons (oscillations) provide a key mechanistic element in most of the brain’s motor, sensory and cognitive operations [1–4]. When sensory input is sampled using overt rhythmic and quasi-rhythmic motor strategies like saccadic search, entrainment of neuronal oscillations enhances the processing of visual information within [5] and beyond the visual system [6] likely improving the transfer of information across a number of brain areas [5,7–12]. Similarly, during covert attentional sampling of rhythmic sensory input stream, entrainment of oscillations to the input rhythm selectively enhances the representation of events in that stream [4,13]. Intriguingly, during covert attention, in lieu of any overt motor behavior or rhythmic sensory cues, visual perceptual threshold appears to fluctuate with the phase of neuronal oscillations [14–20].

While it is clear that these “spontaneous” neuro-behavioral fluctuations depend on a dynamic interplay within the fronto-parietal network in both human [21] and non-human [22] primates, it is unclear whether they are domain-specific or domain-general, and occur outside of the visual system. The results to date concerning the question of parallel effects in auditory processing are inconclusive. Several studies have shown that neuronal oscillations entrain to rhythmic auditory stimulation and that auditory performance co-varies with entrained oscillatory phase [19,23–27]. Neuling et al. [26] observed that detection thresholds depend on the phase of oscillations that were entrained by transcranial stimulation at 10 Hz. In contrast, Ng et al. [27] found that detection of an auditory target in ongoing background noise depends on power and phase in delta/theta but not alpha frequency. Rieke et al. [28] found the TACS-augmentation of endogenous 4 Hz sound-brain phase entrainment accelerates the build-up of auditory streaming. In a gap detection task with an auditory stimulus modulated at 3 Hz, performance clustered around preferred phases rather than being uniformly distributed across time [25]. This phase entrainment has been suggested to play a critical role in speech perception aligning high-excitability auditory phases to speech envelope (for review see [29]). More recently, Ho et al. [30] observed that modulations in perceptual performance oscillate in the left and right ear at different frequencies and phases suggesting that the auditory system might indeed show intrinsic periodicity. Kayser [31] found that complex, temporally extended auditory scenes are sampled rhythmically at the rate of delta frequency range.

A key concern affecting the interpretation of prior studies is that background stimuli often contained energy in several bands including low frequencies [19], making it difficult to determine whether an observed phase concentration is an intrinsic property of a periodically sampling system, or rather, reflects phase locking to the ongoing stimulus rhythm. This concern is further amplified by studies showing a null effect when background stimuli are eliminated; for example, no effect of low frequency phase on auditory perception was observed in quiet (i.e., with no background stimulation; see [19,32]). Similarly, Ilhan et al. [33] report lack of lasting auditory echoes, which was used as evidence for intrinsic perceptual cycling in vision [19]. Consequently, it remains unknown whether auditory perception is continuous with constant threshold or whether it fluctuates with the phase of ongoing EEG between more and less input-favorable states.

We depart from previous studies by using auditory phase-resetting cues followed by a quiet interval interrupted with a monaural target (pitch-discrimination task in quiet) presentation at each subject’s perceptual threshold. Similar procedures have been shown effective in vision [17,18]. We recorded electroencephalography (EEG) to determine whether the phase of prestimulus oscillations predicts accuracy of target discrimination (Figure 1 illustrates the experimental procedure). We tested the predictions that: 1) The distributions of phases between correct and incorrect trials will be centered at different angles as measured with phase bifurcation index (PBI), 2) There will be an inter-individual positive linear relationship between the strength of phase bifurcation indices and behavioral accuracy, and 3) behavioral discrimination performance will undergo periodic modulations rather than showing a uniform distribution across the cue-target time interval. Confirming an effect of pre-stimulus phase on subsequent auditory discrimination in quiet conditions would indicate that the auditory perception performance is not constant but undergoes periodic fluctuations even when there is no entraining rhythm in the sensory stimulus.

**Figure 1.**
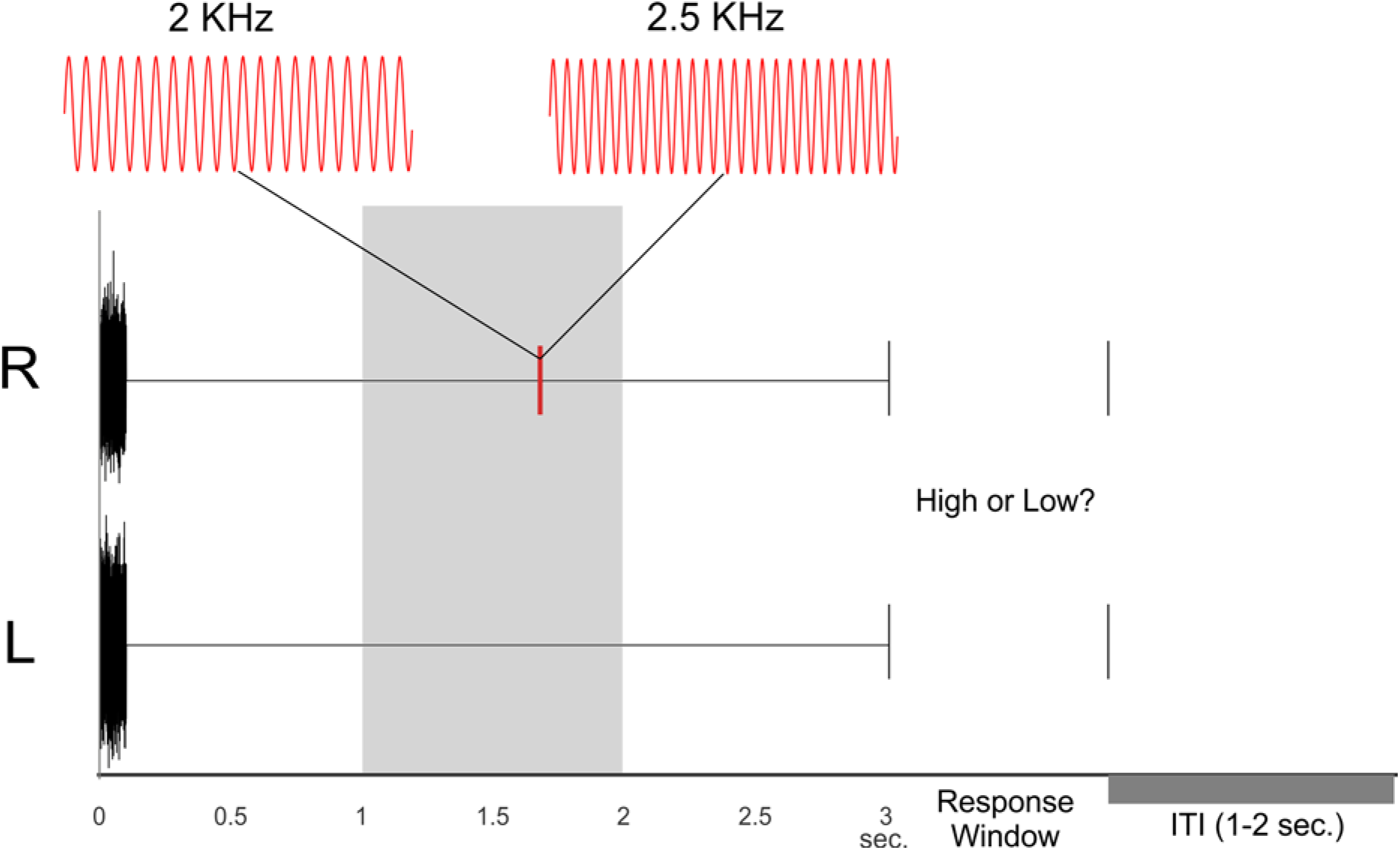
Experimental procedure. Each trail started with a 100 ms long white noise burst followed by a target tone delivered to the left or right ear at a random time point between 1-2 seconds after the onset of the white noise (gray shaded area). Subjects were instructed to decide whether the target was low (2kHz) or high (2.5kHz). Three seconds after the white noise burst, participants were prompted with the response window which required them to indicate if the target was high or low pitch.

## Results

### EEG phase influence on auditory pitch discrimination

The “threshold” pitch discrimination task that we used is diagrammed in Figure1. Overall (average across subjects) auditory pitch discrimination accuracy was 72%. Standard Error of the mean was 0.03. Our goal was to determine if, in lieu of any overt motor behavior or rhythmic sensory cues, the accuracy of auditory pitch discrimination fluctuates with the phase of ongoing neuronal oscillations. To this end, we computed: 1) phase and power consistency for sets of correct and incorrect discrimination trials and 2) a phase bifurcation index (see Methods). A positive phase bifurcation index (PBI) indicates that distributions of oscillatory phases are centered at different angles for correct and incorrect trials. A negative PBI means that only one of the behavioral conditions (correct or incorrect) shows phase concentration while the other condition exhibit random phase across trials.

We compared the spectra of power and PBI as averages across all EEG electrodes for each time point between - 500 ms and 500 ms relative to target onset, for frequencies between 1-30 Hz (see Fig. 2). As expected based on prior work in the visual modality [34,35], we found no significant spectral power differences between correct and incorrect trials (all p > 0.05, FDR corrected).

**Figure 2.**
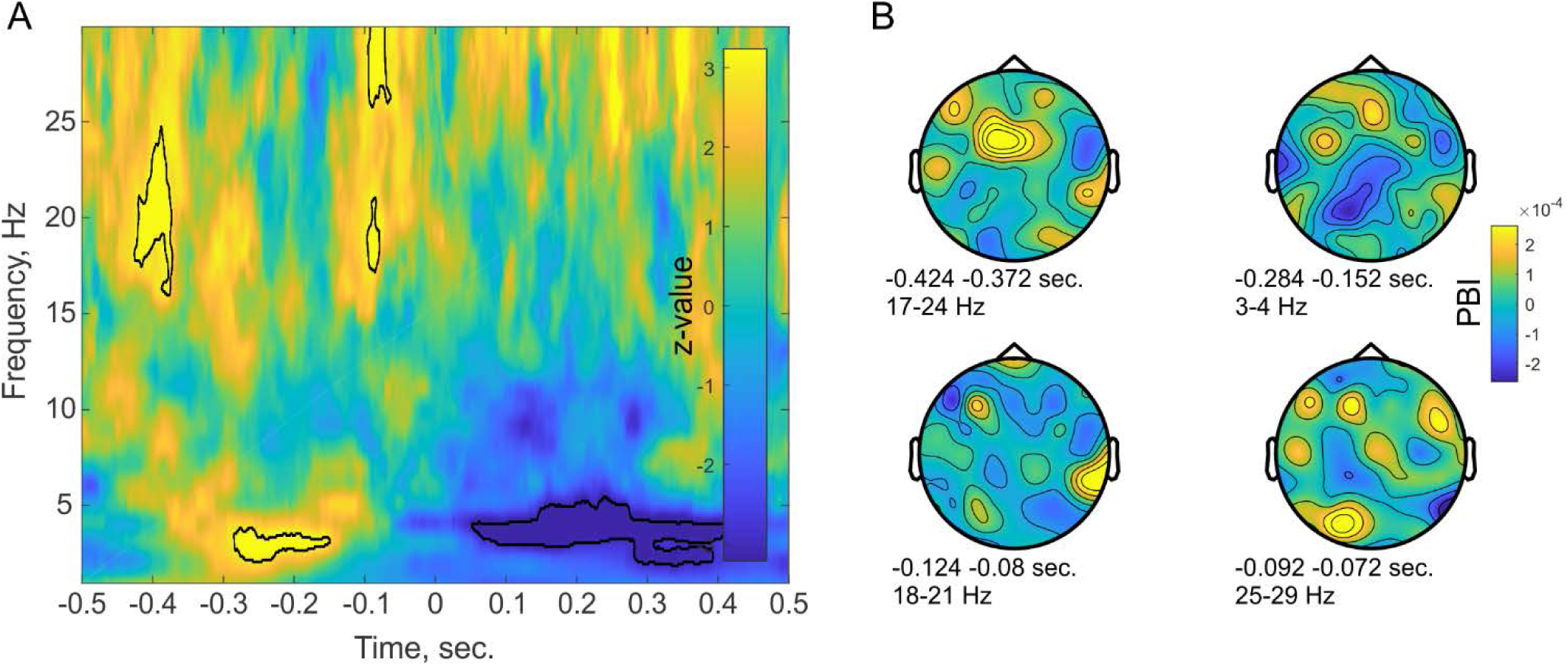
Target-locked Phase Bifurcation Index. A – time-frequency map of PBI averaged across all channels and subjects. X-axis indicates time relative to the onset of the target tone. The color indicates the z-value of the signed-rank test of PBI against 0. Positive values indicate that phase distributions are locked to different phase angles for correct and incorrect trials, while negative values indicate that only one condition is phase locked. Significant clusters are marked with black contour lines. B – topographical plots of the pre-stimulus significant clusters. Times and frequencies of each cluster are indicated in the figure text.

We performed a similar analysis for the phase bifurcation index, testing whether each time-frequency PBI point differed from zero. As shown in Figure 2, we observed four clusters of significantly increased PBI before target onset - one in theta (3-5 Hz) and three in beta (16-29 Hz) frequencies (signed rank test, p < 0.05; FDR corrected for multiple comparisons across all time and frequency points). The strongest effects were observed between - 290 and −150 ms and between −420 to −380 ms for theta and beta frequencies, respectively. Two additional smaller clusters were identified in the beta frequency range at −100 to – 70 ms before stimulus onset. Increased positive PBI indicates that pre-stimulus phase concentration was clustered at different phase angles for correct and incorrect trials; i.e., pre-stimulus phase modifies the likelihood of correctly discriminating between auditory targets.

To ensure there is no inherent phase bias in our data set and that phases were uniformly distributed across all trials, we combined all trials (i.e., correct and incorrect) and used Rayleigh test [36] across all time and frequency points where we observed significant PBI. Importantly, Rayleigh test did not show any departure from uniformity (all p > 0.05, FDR corrected).

Next, we tested whether the transition between phase angles associated with correct and incorrect responses was continuous rather than sudden. We pooled the phase values from the time-frequency points of the PBI clusters into 11 bins (set a priori following the approach of [14]). Next, we calculated the correct to incorrect ratio as a function of the phase bin. We reasoned that if the transition between optimal and non-optimal phases was continuous, it should monotonically change from bins with the highest to the lowest accuracy. Since the exact phase at which performance was highest could vary between subjects, we adjusted each individual’s phase distributions by aligning it to the zero-phase angle for the phase bin at which performance was best. Thus, a trivial feature resulting from the alignment is a peak at zero phase bin observed in Figure 3. Another trivial feature is the difference in accuracy between 0 and +/-pi – we already know from the PBI that correct and incorrect trials cluster at the opposite phase angles. The important non-trivial feature, however, is the monotonic decrease in the ratio of correct and incorrect trials as one moves further from the “optimal” phase. This shows that the phase influence on behavior is continuous - with a small window of sharpest perceptual accuracy monotonically degrading towards the opposite phase angle. To test this monotonic shift in behavioral performance we compared the accuracy values of each phase bin to the adjacent phase bins. We found that most of the differences between adjacent phase bins are significant (t-test, p < 0.05; FDR corrected), indicating a gradual shift in behavioral performance from the ‘optimal’ to the ‘non-optimal’ phase. Importantly, the change in behavioral performance at the shift-point between higher percentage of correct responses to higher percentage of incorrect responses (indicated by the horizontal dashed line in figure 3) was significant in all cases.

**Figure 3.**
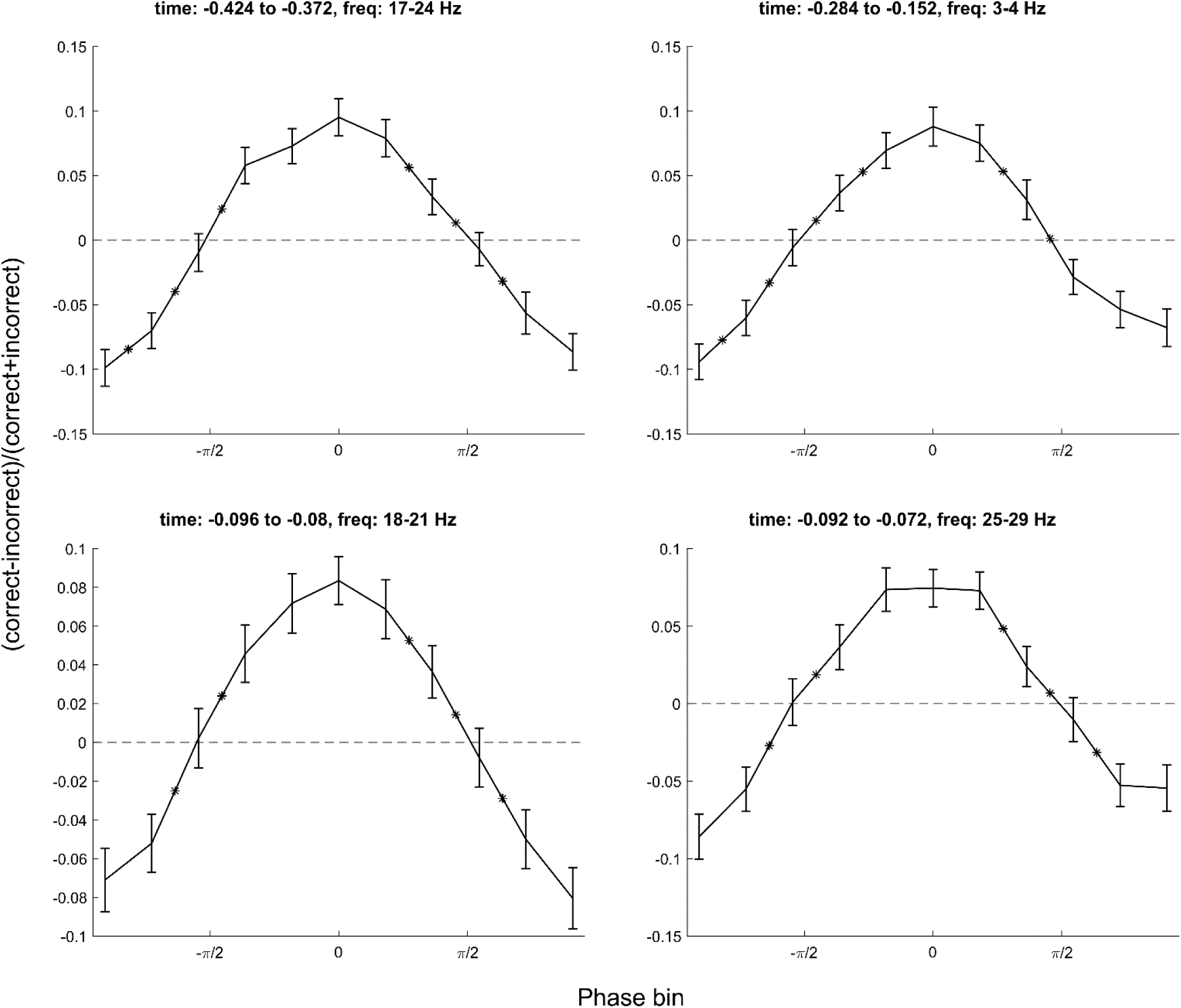
Monotonic relationship between behavioral performance and oscillatory activity. Each panel shows the relationship between spectral phase at each significant time-frequency phase cluster and standardized performance after phases were aligned for each subject such that 0 indicates the optimal phase. Black asterisks indicate a significant change in behavioral performance between adjacent phase bins. Discrimination performance decreases monotonically from the “optimal” phase angles (0 radians) where performance is highest to “non-optimal” phase angles (+/-pi) where performance is lowest.

### Oscillations in behavioral accuracy

To evaluate the assumption that the initial white noise cue resets the ongoing neural oscillations to a common phase angle, we next tested whether accuracy oscillates relative to the cue onset. A target that is presented at random time points after this cue will coincide with different phases of this oscillation. The specific phase at which the target occurs will modulate behavioral performance. To this end, we aggregated individual’s accuracy scores within 20 ms bins relative to the onset of the initial white noise cue. The bin size was selected a priori based on the highest frequency in which we observed a PBI effect in our neural data (see Methods for more details; for a similar approach see [18,30]). To test whether accuracy undergoes rhythmic changes rather than being uniform, we computed the spectrum of the binned behavioral accuracy time course and averaged across participants (Fig. 4). As expected, based on the bifurcation index, the spectrum of accuracy showed the highest peak at 3-4Hz and another peak around 20Hz. To determine whether these peaks are significant, we performed a randomization test in which we shuffled the accuracy bins in time and re-calculated the spectrum of the shuffled accuracy time course. We repeated this procedure 10000 times and set the significance threshold at the 95th percentile of the random distribution. The peak at 3-4 Hz exceeded this statistical threshold and was consistently significant across all tested bin sizes, indicating that behavioral accuracy oscillates at these frequencies. However, the 20 Hz peak was not consistently significant across different phase bins (see Figure S1). Interestingly, the peaks in behavioral accuracy correspond with peaks found with the phase bifurcation index, helping to consolidate our initial observation.

**Figure 4.**
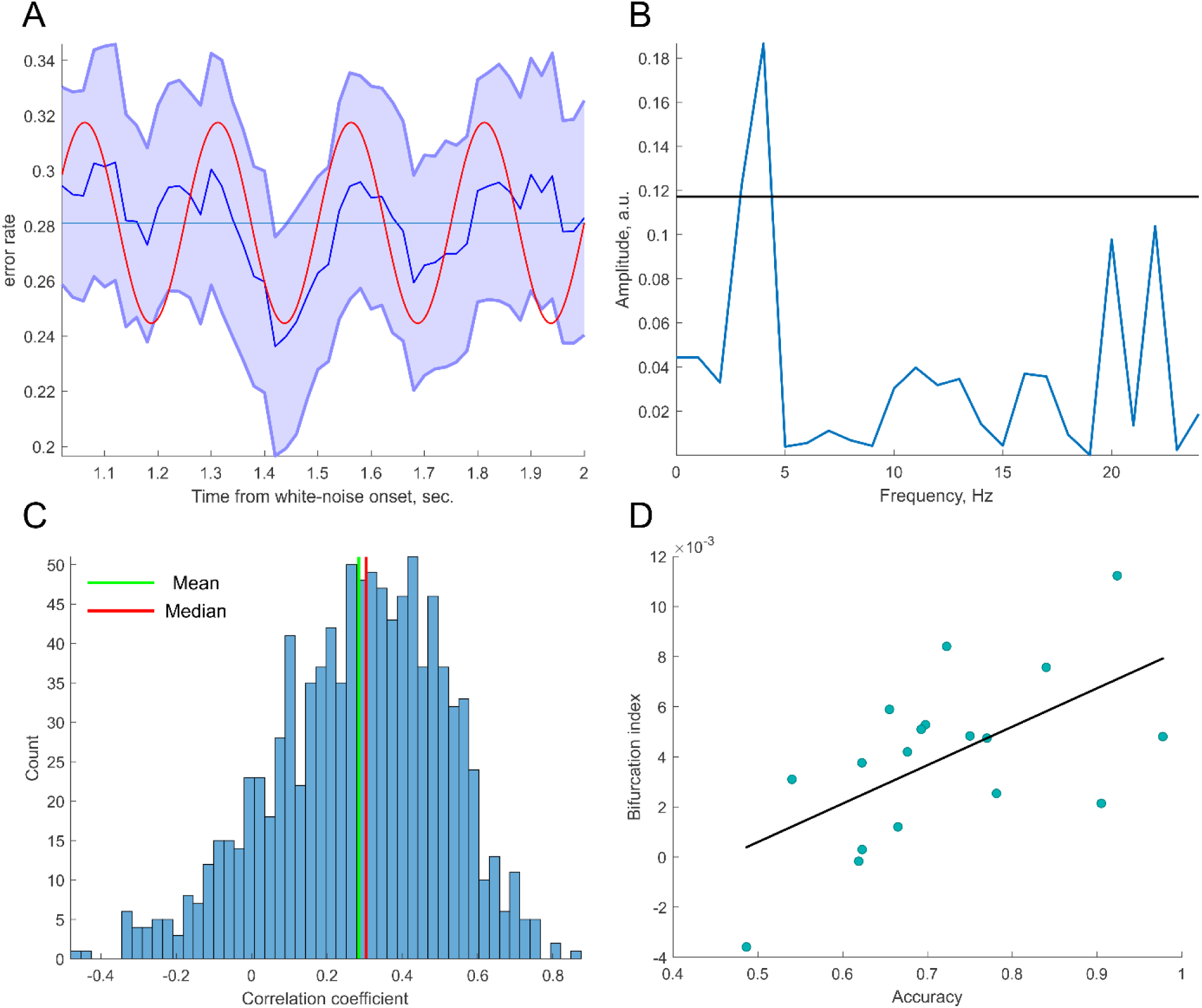
Oscillations in behavioral performance. A – binned behavioral performance as a function of target onset time (blue). Red trace shows a sinusoid at 4 Hz for illustration purposes. Shaded area indicates the SEM. B - spectrum of the binned discrimination accuracy. Black horizontal line indicates the 95th percentile of a random distribution (corresponding to p = 0.05) created by shuffling the behavioral accuracy across time bins. Significant peak was detected at 3-4Hz. C – distribution of correlation coefficients between averaged PBI and behavioral accuracy. The distribution was created by sampling a subset of trials to match the number of trials between subjects (see Methods). D – example correlation between auditory discrimination accuracy throughout the experiment and average bifurcation index around the median correlation value in C. Each point represents a single subject. Subjects that performed better on average also exhibited higher pre-target bifurcation index.

Finally, we tested the inter-individual relation between phase bifurcation index averaged across all significant pre-target clusters and all levels of accuracy during the experiment. Since subjects with higher accuracy had fewer incorrect trials that might bias the PBI, we controlled for the number of trials across participants. We matched the number of trials considered for PBI calculation, by re-sampling a subset of trials for each subject based on the minimum number of incorrect trials across all subjects. We repeated this procedure 1000 times to create a distribution of correlation coefficient values controlled for the number of trials (see Figure 4C). The mean and median of this distribution were ∼0.3 and the distribution was significantly greater than a normal distribution with 0 mean (t-test, p <<.001). Figure 4D shows an example of the correlation between PBI and the accuracy of each subject. This suggests that participants with greater pre-stimulus PBI reached higher accuracy.

## Discussion

A number of studies support the notion of that visual perception is discrete or periodic rather than continuous [14,16–20,37,38][14,16–20,37,38] in that detection/discrimination threshold oscillates with the phase of ongoing EEG or that behavioral performance oscillates as a function of cue-target interval. However, similar intrinsic periodicity has not been demonstrated in the auditory system, despite several attempts to do so. Here, we used an auditory pitch discrimination task with phase resetting cues and monoaural, near-threshold target presentation to address this question. We observed: 1) Phase distributions in the pre-target time-frequency intervals that were centered at different angles for correct and incorrect trials as indexed by positive PBI clusters, 2) Inter-individual positive linear relation between the PBI and behavioral accuracy, and 3) behavioral discrimination performance showing periodic fluctuations at 3-4 Hz (and a trend at 20 Hz). Altogether these findings suggest that auditory perceptual threshold fluctuates with the phase of ongoing neuronal oscillations alternating between optimal and non-optimal sensitivity periods; that is, they support the notion that auditory perception is discrete/periodic rather than continuous.

A series of earlier studies attempted to find evidence for periodic sampling in the auditory system, but were unsuccessful in doing so [19,32,33]. There are two critical changes in the current design that might help understanding the difference between these studies. First, we used a phase resetting cue (i.e., supra-threshold broadband noise lasting for 100 ms). The cue was uninformative about the target timing or its spatial location. Our prediction was that the burst of noise would reset ongoing oscillations to a common phase which we then probe 1-2 sec later with the threshold target presentation. Second, based on findings from [30] that discrimination threshold might oscillate at different rates in the left and right ears, we also used monaural target presentation while previous studies used bilateral targets.

Our findings are consistent with the earlier finding of perceptual fluctuations as a function of prestimulus phase in the visual system [14–16]. Also consistent with previous studies using visual discrimination tasks [34,35], we observed no detectable difference in low frequency power between correct and incorrect trials in our auditory task. In the visual domain, the relationship between pre-stimulus phase and perception was mainly found in the alpha and theta frequency bands [14–20,37]. We also observed an effect in theta frequency (i.e., 3-4 Hz). However, we found additional significant PBI clusters in a higher frequency range (i.e., 16-29 Hz) that were not observed in visual studies. This might reflect an inherent difference between auditory and visual systems. Stimulus presentation at about 40 Hz has been suggested as the optimal frequency for auditory steady-state responses [39,40] compared with lower frequencies observed in the visual system [41].

Previous studies suggest that rhythmic sampling depends on the dynamics of task structure [4,13]. Schroeder and Lakatos proposed that when rhythmic cues are predictive of potential target occurrence, attention operates in a “rhythmic” mode. When task structure is rhythmic [13,42,43] or otherwise predictable [44] low frequency oscillations might be utilized in sensory selection. This “rhythmic” mode entails phase locking to the temporal structure of an attended stream which aligns high-excitability oscillation phases with events in the stream [4]. Consequently, responses in the attended stream are enhanced. In agreement with prior studies in the visual system, the current results indicate that attentive auditory processing operates rhythmically even when the task structure is not rhythmic or predictable. The initial cue in our task alerted the subjects that a target would occur within the next few seconds but did not predict either the precise timing or spatial location (ear) of target presentation. We nevertheless observed that the phase of low frequency oscillations modulated auditory performance.

What might be the source of this periodicity in auditory information processing? Unlike vision or somatosensation, which rely on rhythmic movements to sample the sensory space (e.g., eye movements), there is no obvious rhythmic motoric sampling routine used in auditory processing (but see [45]). Nevertheless, auditory perception is still modulated by top-down inputs from the motor system. Morillon et al. [42] asked their participants to track a slow reference beat with rhythmic finger pressing. They observed that this overt motor activity improved sensitivity to detecting target tones presented in phase with the movement (see also [46]). In a follow up study, Morillon and Baillet [43] used magnetoencephalography to demonstrate that this sensory benefit is driven by top-down influence from the motor cortex. Interestingly, these effects were driven by bursts of beta (18–24 Hz) directed toward auditory regions. Similarly, [47] reported that the motor system modulates the auditory cortex through beta oscillations. Thus, it is possible that the current findings of periodic auditory sampling, particularly in the beta range, reflect ongoing modulations from motor regions.

In summary, the current results show that prestimulus EEG phase differs between correct and incorrect trials in an auditory discrimination task in the absence of a rhythmic (entraining) stimulus. Moreover, we found that discrimination performance oscillates at frequencies similar to those found in concomitant EEG oscillations. These results indicate that, as in vision, auditory perception typically operates in successive periodic cycles, alternating between optimal and non-optimal input processing states.

## Acknowledgments

We thank Kathleen Gao Fan and Jingping Nie for their administrative help, Dr. Luca Iemi and Dr. Saskia Haegens for their helpful comments and discussion.

This work was supported by a grant from the National Institutes of Health, NIDCD, DC014279, P50 MH109429, R01 MH111439 and a grant from The James S. McDonnell Foundation.

## Author contribution

Conceptualization, I.T. and M.L.; Methodology, I.T. and M.L..; Investigation, I.T. and M.L.; Formal Analysis, I.T., Visualization, I.T., Writing – Original Draft, I.T. and M.L.; Writing – Review & Editing, I.T., M.L., N.M. and C.E.S; Funding Acquisition, N.M. and C.E.S.; Resources, N.M. and C.E.S.; Supervision, C.E.S.

## Methods

### RESOURCE AVAILABILITY

#### Lead contact

Further information and requests for resources/code should be directed to the Lead Contact, Idan Tal.

#### Data and code availability

Matlab source code and data generated during this study are available from the Lead Contact on request.

### EXPERIMENTAL MODEL AND SUBJECT DETAILS

#### Subjects

Twenty participants volunteered for the experiment after giving written informed consent and received payment for their participation. All participants had normal or corrected to normal vision and none of them used a hearing aid. Two of the participants were excluded due to unstable behavioral performance during staircase procedure. Eighteen subjects remained in the sample (9 females; mean age: 27 and standard deviation of 7 years; all subjects were right-handed). The experimental procedures were approved by the local IRB at Columbia University.

### METHOD DETAILS

#### Stimuli and procedure

Participants performed an auditory pitch discrimination task between 2 target tones (2kHz and 2.5 kHz) each lasting 10 ms. Each trial started with a 100 ms burst of white noise that was randomly generated on each trial and delivered to both ears. The left and right ear white noise stimuli were time-reversed versions of each other. Then, a target tone was presented monaurally at a random time point between 1-2 seconds following the onset of the white noise. Three seconds after the onset of the white noise, a question appeared on the screen and the subjects had to press a button indicating whether the target tone was high or low in pitch (i.e., 2.5 kHz or 2 kHz respectively; see Figure 1). Inter-trial interval was jittered between 1 and 2 seconds following the participant’s button press.

The experimental session started with a staircase procedure to determine each participant’s individual auditory perceptual threshold. The staircase trial structure was the same as one used in the main experiment. It started with the burst of white noise followed by a pitch discrimination task (see Figure 1). The only difference was target intensity which we decreased following a single correct response and increased when three consecutive incorrect responses were detected (3-down 1-up staircase procedure). This procedure continued until the participant’s performance reached a stable accuracy between 70-75% for a duration of 20 trials. Once their performance stabilized at the desired accuracy we stopped the staircase and calculated the average intensity across the last 20 trials. This individual threshold was then used in the main part of the study. Participants were instructed not to readjust the headphones (Sennheiser CX 3.00) during the experiment to avoid changes in the perceived stimulus intensity. The main experiment consisted of 800 trials (random order - 200 × 2kHz, left ear; 200 × 2kHz, right ear; 200 × 2.5kHz, left ear; 200 × 2.5kHz, right ear).

#### EEG recordings

A Guger Technologies (g.tec) amplifier system was used to record EEG from 62 electrodes mounted in an elastic cap according to the international 10-20 system. Data were recorded in the frequency range from 0.1 to 300 Hz with a sampling rate of 1000 Hz and an online notch filter at 60Hz. Electrode impedances were kept at < 5kΩ.

### QUANTIFICATION AND STATISTICAL ANALYSIS

#### EEG analysis

Data were down sampled off-line to 500 Hz and epoched from 2 seconds before to 2 seconds after target onsets. An automatic artifact rejection excluded epochs in which the signal exceeded 10 standard deviations above the mean of the entire recording period. Eye blinks and eye movements artifacts were removed using ICA as implemented in the FieldTrip toolbox [48]. The remaining data were screened manually for residual artifacts.

The EEG analysis focused on the comparison of pre-stimulus spectral power and phase between correct and incorrect trials. Phase and power were computed by wavelet transform of single trial data for the frequency range of 1-30Hz. The length of the wavelet increased linearly from 3 cycles at 1Hz to 5 cycles at 30Hz to optimize the temporal resolution and stability at lower and higher frequencies respectively. At each time *t* and frequency *f*, the result of the wavelet transform for trial *k* is a complex number in which φ is the phase. The phase concentration across trials is calculated by inter-trial phase coherence (ITC; [49]. We used the following formula to calculate ITC:

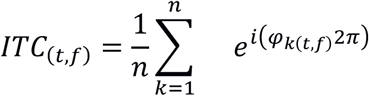

The value of the ITC ranges between 0 and 1, where 0 indicates no synchronization across trials and a value of 1 indicates perfect synchronization. We calculated ITC across correct, incorrect and all trials (correct and incorrect trials pulled together).

We hypothesize that correct and incorrect responses will exhibit pre-target phase concentration at different phase angles. To that end, we used a measure of phase bifurcation index (PBI; [14] that in our case was based on the comparison between ITC for correct and incorrect trials against the ITC calculated across all trials:

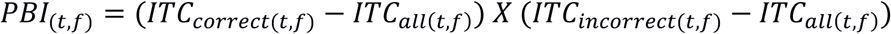

The PBI can take positive or negative values. Importantly, we could use the value of the PBI to infer about the underlying distributions of the phase angles across correct and incorrect trials. For example, if both correct and incorrect conditions have strong ITC centered at the same phase angle – both ITC for correct and incorrect trials will be high. Because correct and incorrect trials are centered at the same phase angle, the ITC for all trials will also be high (see formula 1). Thus, PBI in this case will be ∼0. For the same reason when both conditions have weak ITC (i.e., phase distribution is uniform in both correct and incorrect trials), the ITC for all trials will also show PBI around 0. In turn, when one condition shows high ITC (strong phase concentration at some phase angle) and the other condition has low ITC (uniform phase angle), the ITC all will be higher than the condition with uniform phase but lower than the condition with strong ITC. The PBI in this case will be negative (see formula 2). Finally, when both conditions have strong ITC but centered at distinct phase angles, ITC correct and ITC incorrect will both be larger than ITC all (ITC all will essentially be close to 0 when we combine two distributions centered at distinct phases). The PBI in this case will be positive (for graphical illustration see Figure 1 in [14]). In sum, we could use the value of the PBI to infer about the phase distributions across two conditions. A positive PBI value indicates that both conditions exhibit strong ITC but they are at opposite phases. Negative PBI value indicates that one of the conditions has strong ITC and the other weak ITC. PBI close to 0 indicates one of two cases - both conditions have weak ITC or both conditions have strong ITC at the same phase angles.

To avoid bias in ITC and PBI calculation due to the different number of trials between correct and incorrect responses, we matched the number of trials by selecting the correct trials with the fastest reaction time for each subject assuming that these were more likely to be correct responses as opposed to correct guesses. We computed the PBI for each time and frequency point from −500 to 500 milliseconds and from 1 to 30 Hz around the target onset. Statistical analysis was performed using a signed-rank test for each time and frequency point against 0 and corrected for multiple comparisons using the false discovery rate (FDR) procedure [50].

#### Behavioral analysis

To test whether behavioral performance fluctuates as a function of the target presentation time, we binned the behavioral responses within 20 milliseconds bins and calculated the accuracy in each of these bins. Bin size was determined a priori based on the frequencies of the significant clusters found in the EEG data (20 ms bins allowed us to resolve frequencies up to 25Hz), however, similar results were obtained with different sized bins (ranging from 15 to 40 ms in steps of 5 ms). We then calculated the average of these accuracy curves across subjects and calculated the spectrum of the average curve. Statistical analysis was performed by randomly shuffling the accuracy values across time bins and calculating the spectrum of the shuffled accuracy time course. We repeated this procedure 10000 times and set the significance threshold at the 95th percentile of the random distribution.

## References

1. Bishop, G.H. (1932). Cyclic changes in excitability of the optic pathway of the rabbit. Am. J. Physiol. Content 103, 213–224.

2. Buzsáki, G., and Draguhn, A. (2004). Neuronal oscillations in cortical networks. Science (80-.). 304, 1926–1929.

3. Lakatos, P., Shah, A.S., Knuth, K.H., Ulbert, I., Karmos, G., and Schroeder, C.E. (2005). An oscillatory hierarchy controlling neuronal excitability and stimulus processing in the auditory cortex. J. Neurophysiol. 94, 1904–1911.

4. Schroeder, C.E., and Lakatos, P. (2009). Low-frequency neuronal oscillations as instruments of sensory selection. Trends Neurosci. 32, 9–18.

5. Barczak, A., Haegens, S., Ross, D.A., McGinnis, T., Lakatos, P., and Schroeder, C.E. (2019). Dynamic modulation of cortical excitability during visual active sensing. Cell Rep. 27, 3447–3459.

6. Leszczynski, M., Staudigl, T., Chaieb, L., Enkirch, S.J., Fell, J., and Schroeder, C.E. (2020). Saccadic modulation of neural activity in the human anterior thalamus during visual active sensing. BioRxiv.

7. Purpura, K.P., Kalik, S.F., and Schiff, N.D. (2003). Analysis of perisaccadic field potentials in the occipitotemporal pathway during active vision. J. Neurophysiol. 90, 3455–3478.

8. Rajkai, C., Lakatos, P., Chen, C.-M., Pincze, Z., Karmos, G., and Schroeder, C.E. (2008). Transient cortical excitation at the onset of visual fixation. Cereb. Cortex 18, 200–209.

9. Bartlett, A.M., Ovaysikia, S., Logothetis, N.K., and Hoffman, K.L. (2011). Saccades during object viewing modulate oscillatory phase in the superior temporal sulcus. J. Neurosci. 31, 18423–18432.

10. Ito, J., Maldonado, P., Singer, W., and Grün, S. (2011). Saccade-related modulations of neuronal excitability support synchrony of visually elicited spikes. Cereb. Cortex 21, 2482–2497.

11. Jutras, M.J., Fries, P., and Buffalo, E.A. (2013). Oscillatory activity in the monkey hippocampus during visual exploration and memory formation. Proc. Natl. Acad. Sci. 110, 13144–13149.

12. Leszczynski, M., and Schroeder, C.E. (2019). The Role of Neuronal Oscillations in Visual Active Sensing. Front. Integr. Neurosci. 13, 1–9.

13. Lakatos, P., Karmos, G., Mehta, A.D., Ulbert, I., and Schroeder, C.E. (2008). Entrainment of neuronal oscillations as a mechanism of attentional selection. Science (80-.). 320, 110–113. Available at: http://www.ncbi.nlm.nih.gov/pubmed/18388295.

14. Busch, N.A., Dubois, J., and VanRullen, R. (2009). The Phase of Ongoing EEG Oscillations Predicts Visual Perception. J. Neurosci. 29, 7869–7876. Available at: http://www.jneurosci.org/cgi/doi/10.1523/JNEUROSCI.0113-09.2009.

15. Busch, N.A., and VanRullen, R. (2010). Spontaneous EEG oscillations reveal periodic sampling of visual attention. Proc. Natl. Acad. Sci. U. S. A. 107, 16048–16053.

16. Mathewson, K.E., Fabiani, M., Gratton, G., Beck, D.M., and Lleras, A. (2010). Rescuing stimuli from invisibility: Inducing a momentary release from visual masking with pre-target entrainment. Cognition 115, 186–191. Available at: http://dx.doi.org/10.1016/j.cognition.2009.11.010.

17. Landau, A.N., and Fries, P. (2012). Attention samples stimuli rhythmically. Curr. Biol. 22, 1000–1004. Available at: http://dx.doi.org/10.1016/j.cub.2012.03.054.

18. Fiebelkorn, I.C., Saalmann, Y.B., and Kastner, S. (2013). Rhythmic sampling within and between objects despite sustained attention at a cued location. Curr. Biol. 23, 2553–2558. Available at: http://dx.doi.org/10.1016/j.cub.2013.10.063.

19. VanRullen, R., Zoefel, B., and Ilhan, B. (2014). On the cyclic nature of perception in vision versus audition. Philos. Trans. R. Soc. B Biol. Sci. 369.

20. VanRullen, R. (2016). Perceptual Cycles. Trends Cogn. Sci. 20, 723–735. Available at: http://dx.doi.org/10.1016/j.tics.2016.07.006.

21. Helfrich, R.F., Fiebelkorn, I.C., Szczepanski, S.M., Lin, J.J., Parvizi, J., Knight, R.T., Helfrich, R.F., Fiebelkorn, I.C., Szczepanski, S.M., Lin, J.J., et al. (2018). Neural Mechanisms. Neuron 99, 854-865.e5. Available at: https://doi.org/10.1016/j.neuron.2018.07.032

22. Fiebelkorn, I.C., Pinsk, M.A., and Kastner, S. (2018). Article A Dynamic Interplay within the Frontoparietal A Dynamic Interplay within the Frontoparietal Network Underlies Rhythmic Spatial Attention. Neuron 99, 842-853.e8. Available at: https://doi.org/10.1016/j.neuron.2018.07.038

23. Lakatos, P., Musacchia, G., O’Connell, M.N., Falchier, A.Y., Javitt, D.C., and Schroeder, C.E. (2013). The spectrotemporal filter mechanism of auditory selective attention. Neuron 77, 750–761.

24. Zion Golumbic, E.M., Ding, N., Bickel, S., Lakatos, P., Schevon, C.A., McKhann, G.M., Goodman, R.R., Emerson, R., Mehta, A.D., Simon, J.Z., et al. (2013). Mechanisms underlying selective neuronal tracking of attended speech at a “cocktail party.” Neuron 77, 980–991. Available at: http://dx.doi.org/10.1016/j.neuron.2012.12.037.

25. Henry, M.J., and Obleser, J. (2012). Frequency modulation entrains slow neural oscillations and optimizes human listening behavior. Proc. Natl. Acad. Sci. U. S. A. 109, 20095–20100.

26. Neuling, T., Rach, S., Wagner, S., Wolters, C.H., and Herrmann, C.S. (2012). Good vibrations: Oscillatory phase shapes perception. Neuroimage 63, 771–778. Available at: http://dx.doi.org/10.1016/j.neuroimage.2012.07.024.

27. Ng, B.S.W., Schroeder, T., and Kayser, C. (2012). A precluding but not ensuring role of entrained low-frequency oscillations for auditory perception. J. Neurosci. 32, 12268–12276.

28. Riecke, L., Sack, A.T., and Schroeder, C.E. (2015). Endogenous delta/theta sound-brain phase entrainment accelerates the buildup of auditory streaming. Curr. Biol. 25, 3196–3201.

29. Zoefel, B., and VanRullen, R. (2015). The Role of High-Level Processes for Oscillatory Phase Entrainment to Speech Sound. Front. Hum. Neurosci. 9, 1–12. Available at: http://journal.frontiersin.org/Article/10.3389/fnhum.2015.00651/abstract.

30. Ho, H.T., Leung, J., Burr, D.C., Alais, D., and Morrone, M.C. (2017). Auditory Sensitivity and Decision Criteria Oscillate at Different Frequencies Separately for the Two Ears. Curr. Biol., 1–7. Available at: http://linkinghub.elsevier.com/retrieve/pii/S0960982217313209 https://m.medicalxpress.com/news/2017-11-neuroscience-evidence-brain-strobing-constant.html.

31. Kayser, C. (2019). Evidence for the rhythmic perceptual sampling of auditory scenes. Front. Hum. Neurosci. 13, 249.

32. Zoefel, B., and Heil, P. (2013). Detection of near-threshold sounds is independent of eeg phase in common frequency bands. Front. Psychol. 4, 1–17.

33. Ilhan, B., and VanRullen, R. (2012). No Counterpart of Visual Perceptual Echoes in the Auditory System. PLoS One 7.

34. Iemi, L., Chaumon, M., Crouzet, S.M., and Busch, N.A. (2017). Spontaneous neural oscillations bias perception by modulating baseline excitability. J. Neurosci. 37, 807–819.

35. Samaha, J., Gosseries, O., and Postle, B.R. (2017). Distinct oscillatory frequencies underlie excitability of human occipital and parietal cortex. J. Neurosci. 37, 2824–2833.

36. Mardia, K. V (1975). Statistics of Directional Data. J. R. Stat. Soc. Ser. B 37, 349–393. Available at: http://www.jstor.org/stable/2984782.

37. VanRullen, R., and Koch, C. (2003). Is perception discrete or continuous? Trends Cogn. Sci. 7, 207–213. Available at: https://doi.org/10.1016/S1364-6613(03)00095-0

38. Fiebelkorn, I.C., and Kastner, S. (2019). A Rhythmic Theory of Attention. Trends Cogn. Sci. 23, 87–101. Available at: https://doi.org/10.1016/j.tics.2018.11.009

39. Galambos, R., Makeig, S., and Talmachoff, P.J. (1981). A 40-Hz auditory potential recorded from the human scalp. Proc. Natl. Acad. Sci. U. S. A. 78, 2643–2647.

40. Stapellst, D.R., Linden, D., Suffield, J.B., Hamel, G., and Picton, T.W. (1984). Human auditory steady state potentials. Ear Hear. 5, 105–113.

41. Ding, J., Sperling, G., and Srinivasan, R. (2006). Attentional modulation of SSVEP power depends on the network tagged by the flicker frequency. Cereb. Cortex 16, 1016–1029. Available at: https://www.ncbi.nlm.nih.gov/pubmed/16221931.

42. Morillon, B., Schroeder, C.E., and Wyart, V. (2014). Motor contributions to the temporal precision of auditory attention. Nat. Commun. 5, 1–9. Available at: http://dx.doi.org/10.1038/ncomms6255 http://www.pubmedcentral.nih.gov/articlerender.fcgi?artid=4199392&tool=pmcentrez&rendertype=abstract.

43. Morillon, B., and Baillet, S. (2017). Motor origin of temporal predictions in auditory attention. Proc. Natl. Acad. Sci., 201705373. Available at: http://www.pnas.org/lookup/doi/10.1073/pnas.1705373114.

44. Morillon, B., Schroeder, C.E., Wyart, V., and Arnal, L.H. (2016). Temporal prediction in lieu of periodic stimulation. J. Neurosci. 36, 2342–2347.

45. Gruters, K.G., Murphy, D.L.K., Jenson, C.D., Smith, D.W., Shera, C.A., and Groh, J.M. (2018). The eardrums move when the eyes move: A multisensory effect on the mechanics of hearing. Proc. Natl. Acad. Sci. 115, E1309–E1318.

46. Zalta, A., Petkoski, S., and Morillon, B. (2020). Natural rhythms of periodic temporal attention. Nat. Commun. 11, 1–12.

47. Abbasi, O., and Gross, J. (2019). Beta-band oscillations play an essential role in motor–auditory interactions. Hum. Brain Mapp., 1–10.

48. Oostenveld, R., Fries, P., Maris, E., and Schoffelen, J.-M.M. (2010). FieldTrip: open source software for advanced analysis of MEG, EEG, and invasive electrophysiological data. Comput. Intell. Neurosci. 2011.

49. Lachaux, J.-P., Rodriguez, E., Martinerie, J., and Varela, F.J. (1999). Measuring phase synchrony in brain signals. Hum. Brain Mapp. 8, 194–208. Available at: https://doi.org/10.1002/(SICI)1097-0193(1999)8:4%3C194::AID-HBM4%3E3.0.CO

50. Benjamini, Y., Hochberg, Y., and Benjamini, Yoav H.Y. (1995). Controlling the False Discovery Rate: A Practical and Powerful Approach to Multiple Testing. J. R. Stat. Soc. Ser. B 57, 289–300. Available at: http://www.stat.purdue.edu/~doerge/BIOINFORM.D/FALL06/Benjamini and Y FDR.pdf http://engr.case.edu/ray_soumya/mlrg/controlling_fdr_benjamini95.pdf.

