## Supplemental Figure 1 for "Does the phase of ongoing EEG oscillations predict auditory perception?"

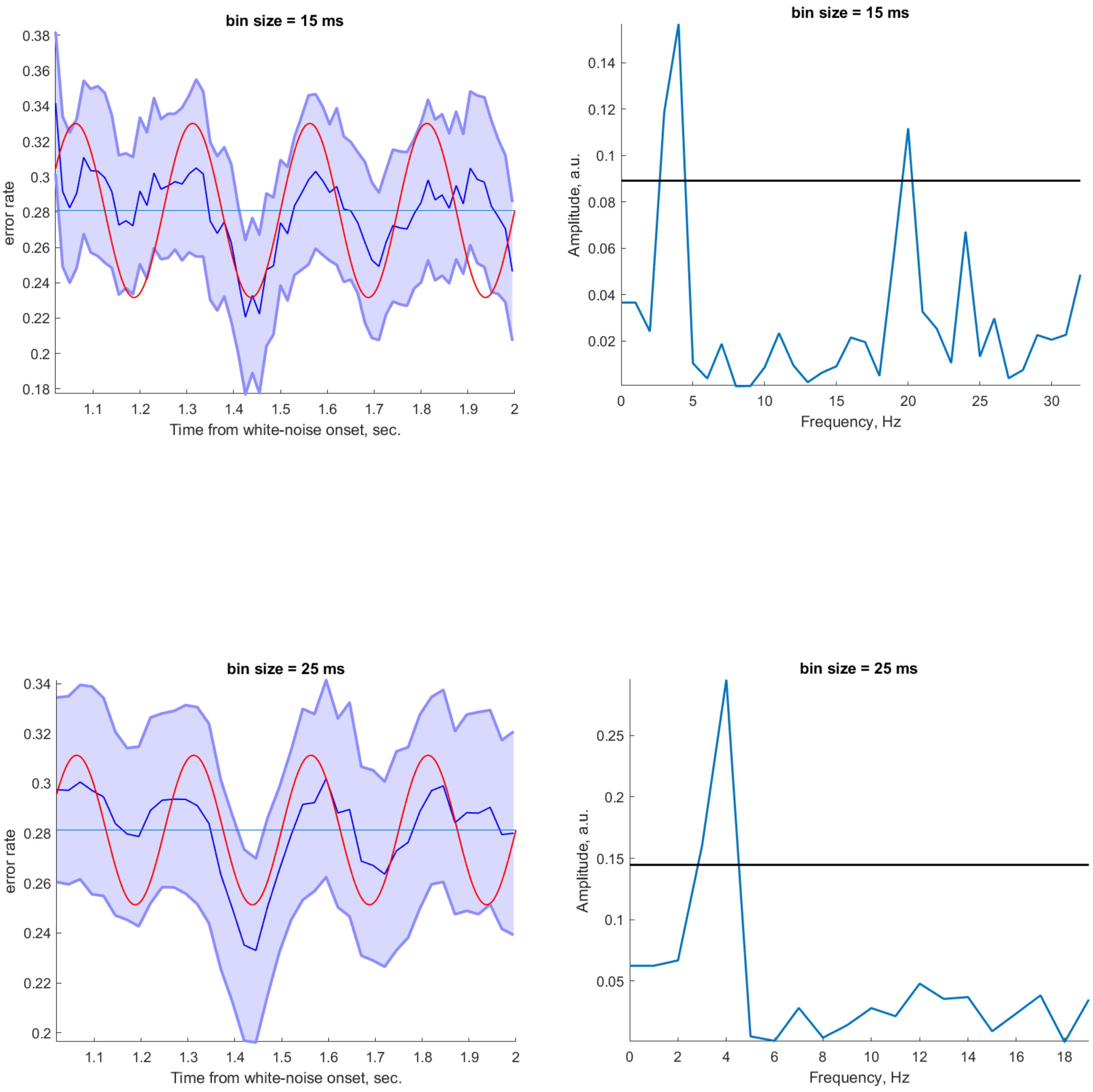

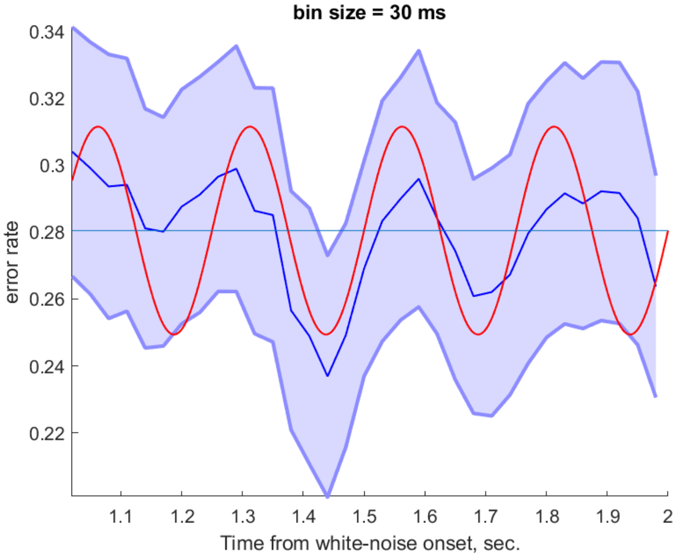
bin size = 30 ms
error rate
0.34
0.32
0.3
0.28
0.26
0.24
0.22
1.1
1.2
1.3
1.4
1.5
1.6
1.7
1.8
1.9
2
Time from white-noise onset, sec.

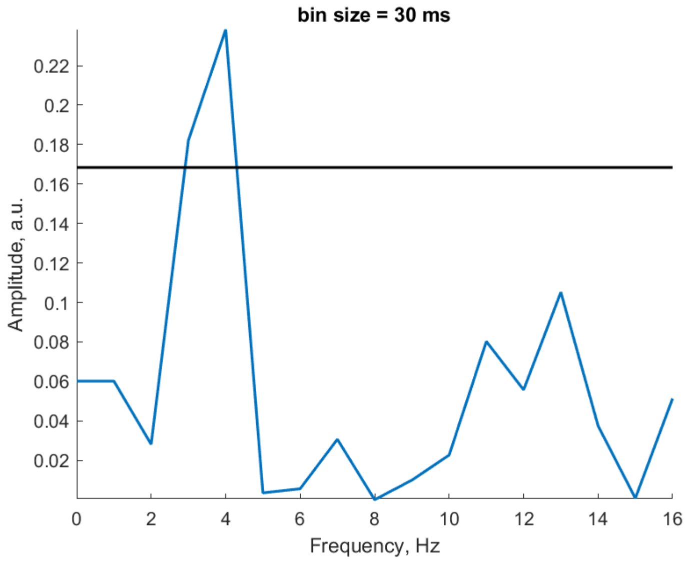
bin size = 30 ms
Amplitude, a.u.
0.22
0.2
0.18
0.16
0.14
0.12
0.1
0.08
0.06
0.04
0.02
0
2
4
6
8
10
12
14
16
Frequency, Hz

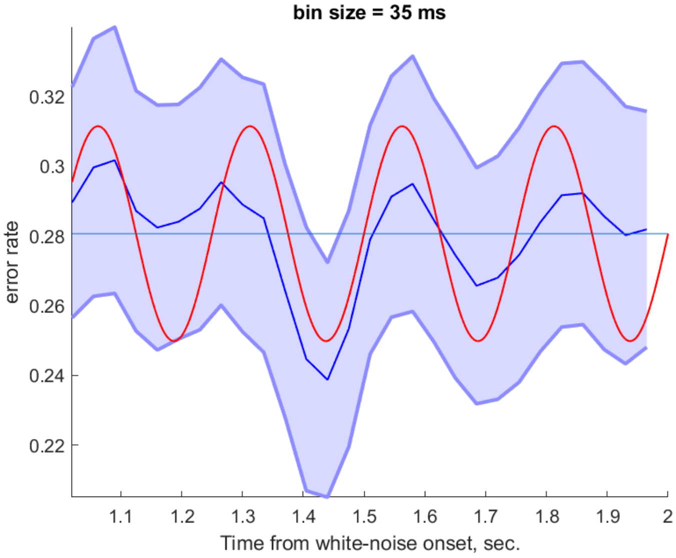
bin size = 35 ms
error rate
0.32
0.3
0.28
0.26
0.24
0.22
1.1
1.2
1.3
1.4
1.5
1.6
1.7
1.8
1.9
2
Time from white-noise onset, sec.

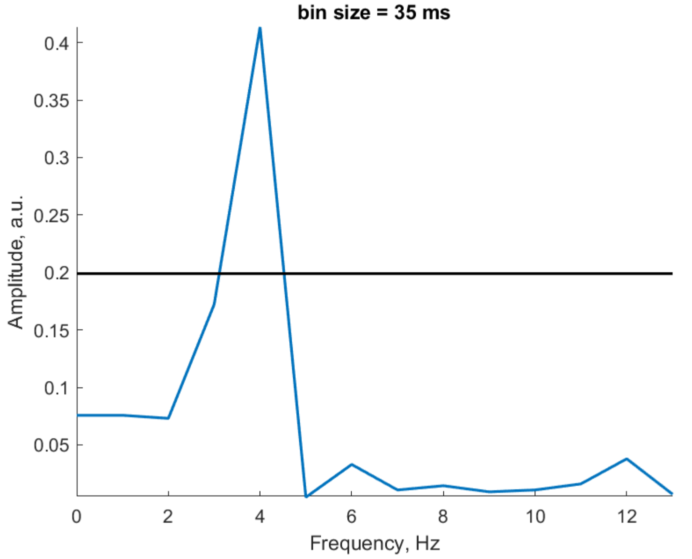
bin size = 35 ms
Amplitude, a.u.
0.4
0.35
0.3
0.25
0.2
0.15
0.1
0.05
0
2
4
6
8
10
12
Frequency, Hz

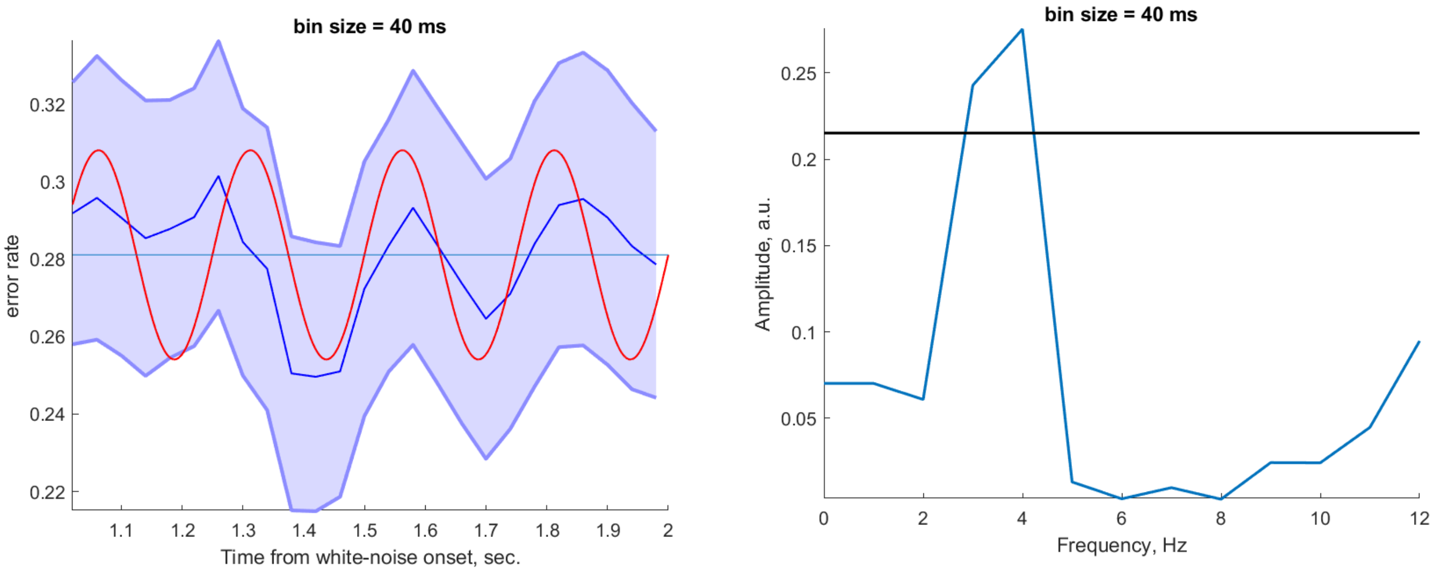


***Figure S1.*** *Oscillations in behavioral performance for different bin sizes. Left - binned behavioral performance as a function of target onset time (blue). Red trace shows a sinusoid at 4 Hz for illustration purposes. Shaded area indicates the SEM. Right - spectrum of the binned discrimination accuracy. Black horizontal line indicates the 95th percentile of a random distribution (corresponding to p = 0.05) created by shuffling the behavioral accuracy across time bins. Different bin sizes are indicated in the title of each panel. Significant peaks were consistently detected at 3-4Hz. Another peak at 20Hz was detected for a bin size of 15 ms and was not consistent across different bin sizes (note that for longer bin sizes the sampling rate was too low to resolve 20Hz).*
